## Supplementary Material for "‘Identification of RING E3 pseudoligases in the TRIM protein family’"

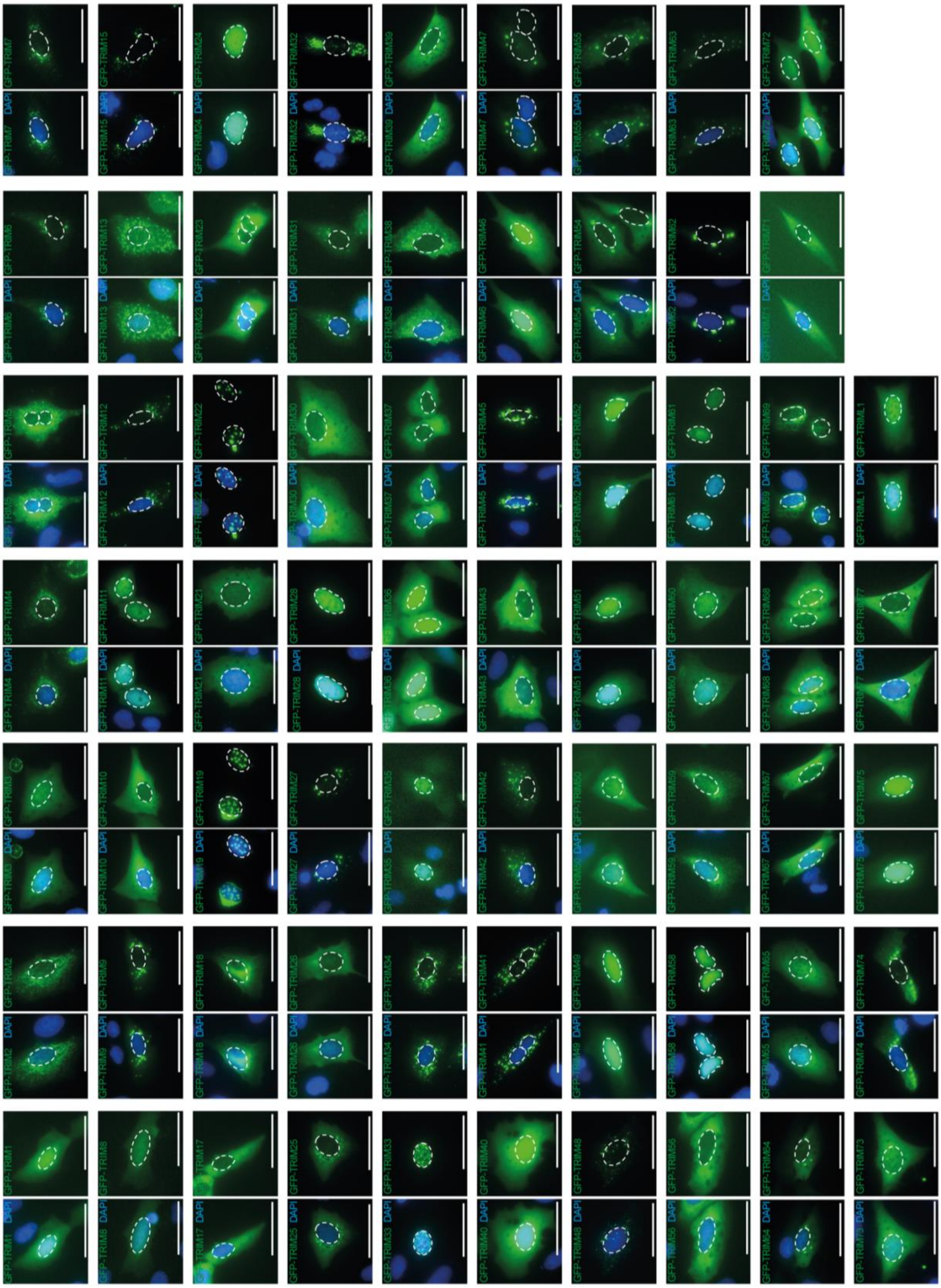

**Supplementary Figure 1 – Localisation of RING-containing TRIM proteins**

Representative images U2OS cells transiently over-expressing GFP-tagged RING-containing TRIM proteins 1 through to L1 (green), which were fixed and stained with DAPI (blue). Dashed lines denote the nucleus and the scale bars represent 50 μm. Images represent n≥3.

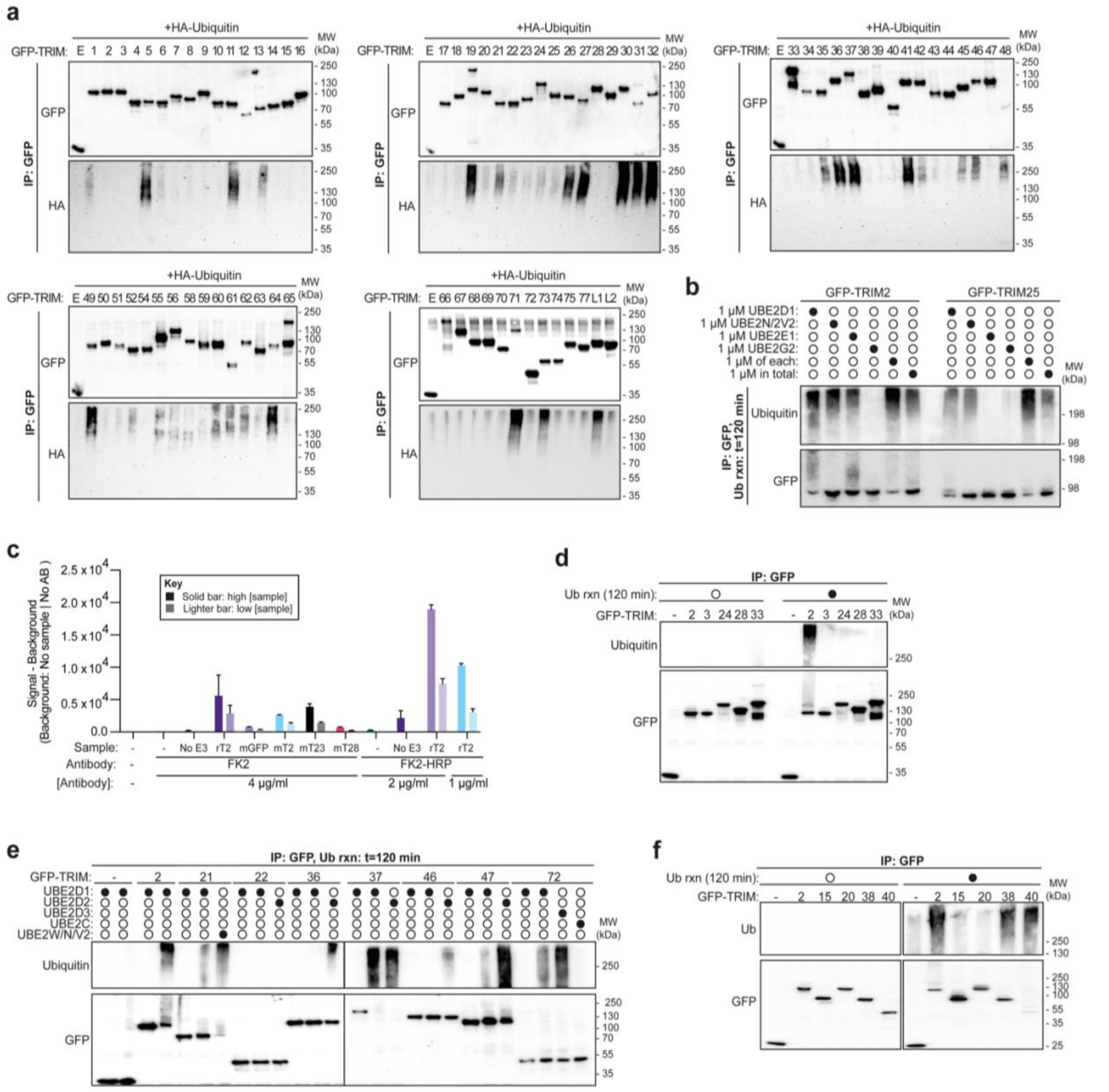

**Supplementary Figure 2 – Data supporting screening TRIM family for in-cell and *in vitro* auto-ubiquitination activities**

**a**, Relating to Figure 1f, representative western blots using the indicated antibodies to detect in-cell TRIM ubiquitination using immunoprecipitated GFP-TRIMs transiently co-over-expressed with HA-Ubiquitin in HEK293T cells, treated with 10 μM PR619 and 10 μM MG132 4 h before lysis. **b**, Western blots using the indicated antibodies, demonstrating the efficacy of mixing E2 enzymes for *in vitro* auto-ubiquitination assays with GFP-tagged TRIM2 and TRIM25 isolated from HEK293T cells (n=2). **c**, Graph representing the validation of an ELISA assay specific to conjugated ubiquitin for assessment of *in vitro* auto-ubiquitination, using reaction mix without E3 or GFP only (negative controls), recombinant TRIM2 RING domain (rT2: positive control) and full-length mammalian cell-derived TRIMs (mT2, mT23, or mT28) at high or low dilutions in the assay. FK2 antibody followed by anti-mouse-HRP was used to trial an indirect ELISA, whilst FK2-HRP was able to demonstrate the enhanced sensitivity of a direct ELISA in this case. Error bars represent mean of duplicates ± SEM. **d**, Western blot using the indicated antibodies against immunoprecipitated GFP-tagged TRIM proteins from HEK293T cells and then used in an *in vitro* ubiquitination assay for 0 or 120 min with a mix of E2 enzymes as in Figure 1f (n=3). **e**, Western blot using the indicated antibodies against immunoprecipitated GFP-tagged TRIM proteins purified from HEK293T cells, which were then used in an *in vitro* ubiquitination assay for 0 or 120 min with different E2 enzymes (UBE2C, UBE2D1, UBE2D2, UBE2D3, or a mix of UBE2N/UBE2V2/UBE2W) (n=3). **f**, Western blot using the indicated antibodies against immunoprecipitated GFP-tagged TRIM proteins purified from HEK293T cells, which were then used in an *in vitro* ubiquitination assay for 0 or 120 min with the mix of E2s as specified in Fig. 1f (n=3).

| TRIM class | Class members | RING overview AlphaFold2 | Core RING AlphaFold2 | Notable predicted features |
|---|---|---|---|---|
| Class I | TRIM1<br>TRIM9<br>TRIM18<br>TRIM36<br>TRIM46<br>TRIM67 | 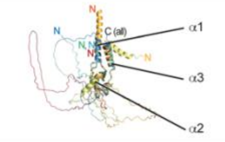 | 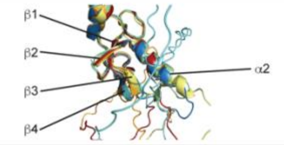 | TRIM9, TRIM36, TRIM46, TRIM67<br>Long loop between α2 and β3 |
| Class II | TRIM54<br>TRIM55<br>TRIM63 | 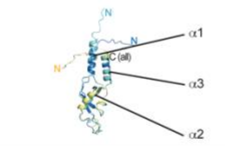 | 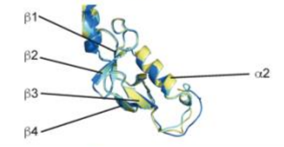 | |
| Class III | TRIM42 | 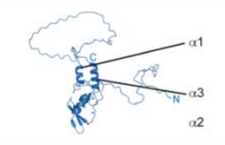 | 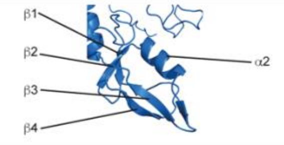 | TRIM42<br>Long N-terminus preceding RING domain |
| Class IV | TRIM4 TRIM26 TRIM50<br>TRIM5* TRIM27 TRIM51<br>TRIM6 Trim30 TRIM58<br>TRIM7 TRIM34 TRIM60<br>TRIM10 TRIM35 TRIM62<br>TRIM11 TRIM38 TRIM64<br>Trim12 TRIM39 TRIM65<br>TRIM15 TRIM41 TRIM68<br>TRIM17 TRIM43 TRIM69*<br>TRIM21* TRIM47 TRIM72<br>TRIM22 TRIM48 TRIM75<br>TRIM25* TRIM49 TRIM77<br>TRIML1 | 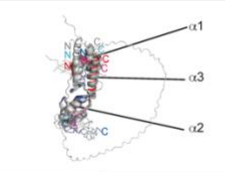 | 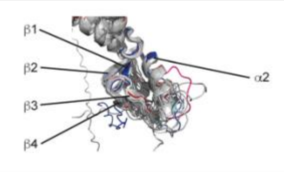 | TRIM6<br>Unfolded α2<br>TRIM15<br>Unfolded α1, 2, 3 and β3, 4<br>TRIM51<br>Unfolded β3, 4<br>TRIM41<br>Long loop between α2 and β3<br>See Fig. 2 |
| Class V | TRIM8<br>TRIM19*<br>TRIM31<br>TRIM40<br>TRIM52<br>TRIM56*<br>TRIM61<br>TRIM73<br>TRIM74 | 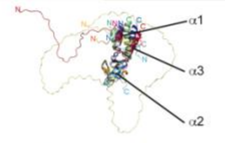 | 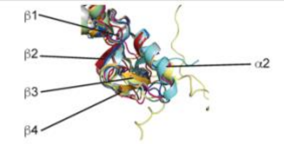 | TRIM8, TRIM19, TRIM40<br>Unfolded β3, 4<br>TRIM52<br>Long loop between α2 and β3<br>TRIM19<br>Long N-terminus preceding RING domain |
| Class VI | TRIM24<br>TRIM28*<br>TRIM33 | 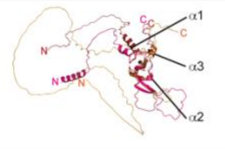 | 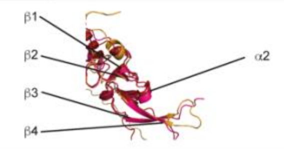 | TRIM24, TRIM28<br>Largely unfolded α1, 2, 3<br>TRIM33<br>Unfolded α1, largely unfolded α2, 3<br>All<br>Long N-terminus preceding RING domain |
| Class VII | TRIM2*<br>TRIM3<br>TRIM32*<br>TRIM71 | 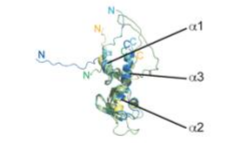 | 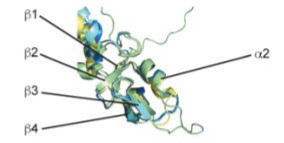 | TRIM3<br>Unfolded α1 |
| Class VIII | TRIM37* | 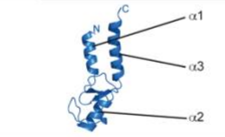 | 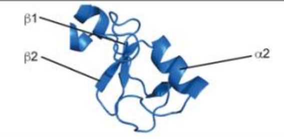 | TRIM37<br>Unfolded β3, 4 |
| Class IX | TRIM23* | 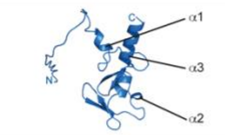 | 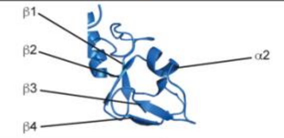 | |
| Class X | TRIM45 | 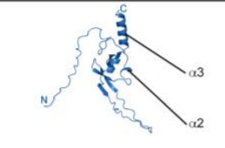 | 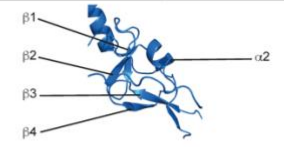 | TRIM45<br>Unfolded α1, long loop between α2 and β3 |
| Class XI | TRIM13<br>TRIM59 | 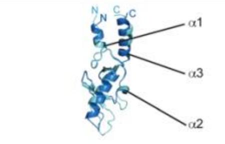 | 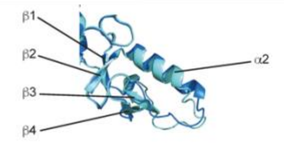 | TRIM59<br>Unfolded β3, 4 |

**Supplementary Figure 3 – *in silico* analyses of TRIM family proteins highlights notable structural differences, including in TRIMs identified as inactive**

Table of AlphaFold2 structural predictions of TRIM RING domains (N-terminal Met to end of $\alpha 3$ helix, or with a sequence of corresponding length where no $\alpha 3$ helix is predicted), aligned and arranged according to class designation, with descriptions of notably divergent structural features given for some TRIMs. * indicates TRIMs with a RING domain structure available in the PDB (TRIM2: 7ZJ3; TRIM5: 4TKP; TRIM19: 5YUF; TRIM21: 5OLM, 6FGA, 7BBD, 6S53; TRIM23: 5CZV; TRIM25: 5EYA, 5FER; TRIM28: 6QAJ, 6I9H; TRIM32: 5FEY; TRIM37: 3LRQ; TRIM56: 5JW7; TRIM69: 6YXE)

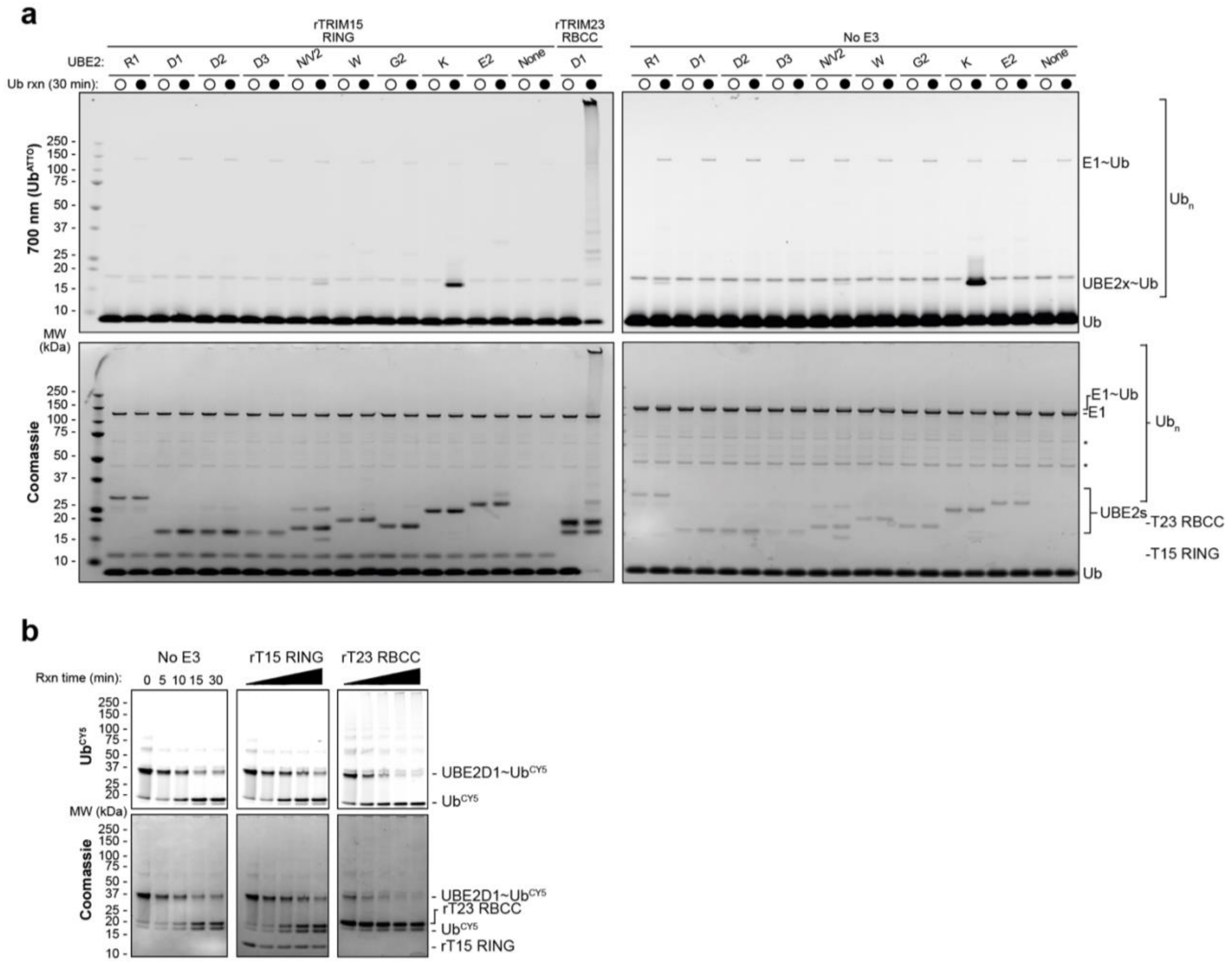

**Supplementary Figure 4 – Data supporting TRIM15 has a monomeric RING domain that does not demonstrate ubiquitin ligase activity in isolation**

**a**, Fluorescent and Coomassie gel scans of auto-ubiquitination reaction carried out at 30 °C 30 min using 1 μM of each of the indicated E2 enzymes (UBE2R1, D1, D2, D3, N/V2, W, G2, K, or E2) with 1 μM UBA1, 50 μM ubiquitin, 1 μM $Ub^{ATTO}$, and 3 mM ATP, using no E3 or 4 μM of recombinant TRIM23 RBCC or TRIM15 RING (n=3). **b**, Fluorescent and Coomassie gel scans showing the discharge of $Ub^{CY5}$ from pre-charged UBE2D1 onto free lysine in solution, with 4 μM recombinant TRIM23 RBCC domain or recombinant TRIM15 RING domain, or with no E3 (see Fig. 3b).

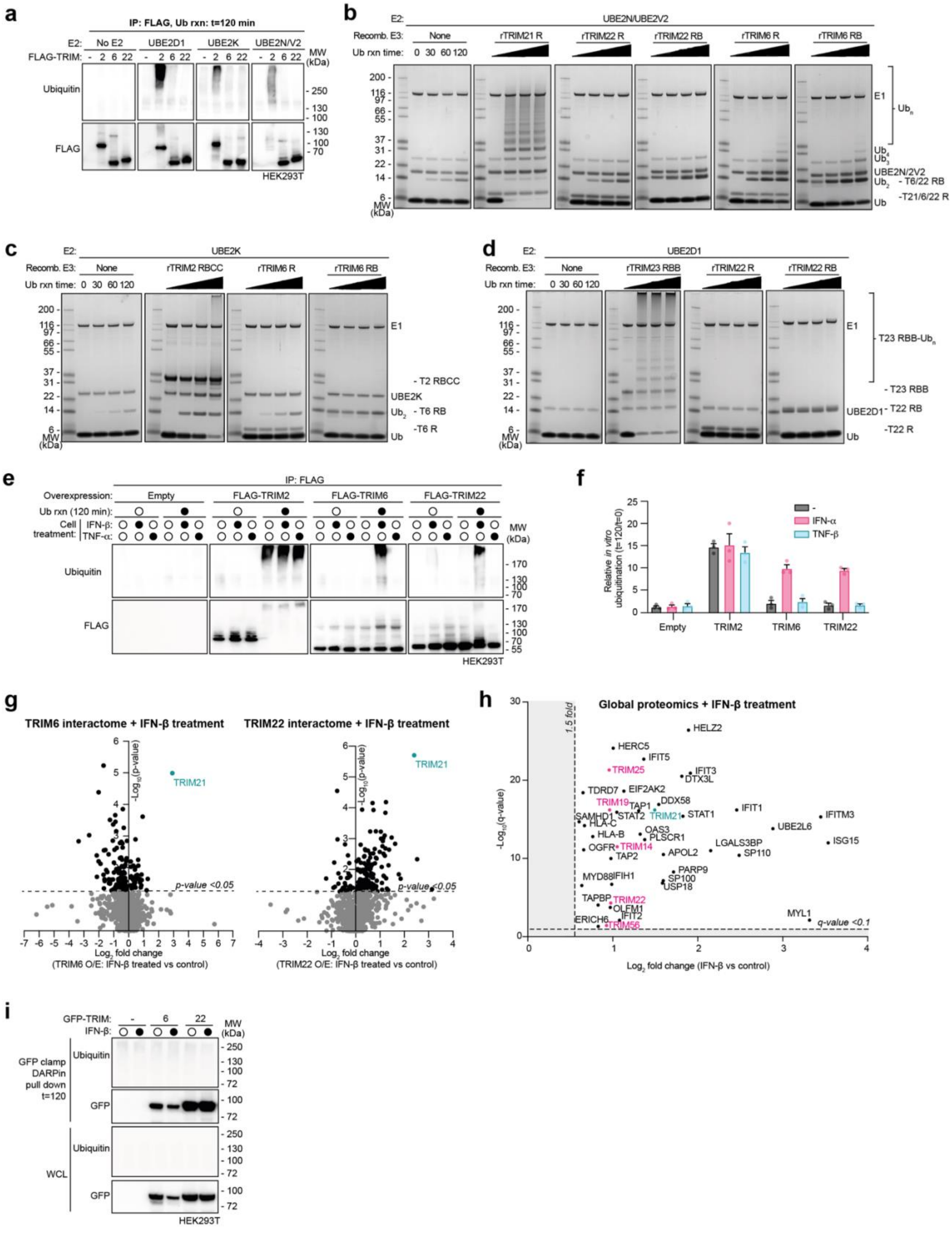

**Supplementary Figure 5 – Analysis of TRIM6 and TRIM22 activity in response to interferon signalling**

**a**, Western blots using the indicated antibodies against immunoprecipitated FLAG-tagged TRIM proteins purified from HEK293T cells, which were then used in an *in vitro* ubiquitination assay for 120 min with 1 μM of specified E2 enzymes (or no E2 control) with 1 μM UBA1, 50 μM ubiquitin, and 3 mM ATP (n=3). **b-d**, Coomassie gels of *in vitro* auto-ubiquitination assays for the indicated times as described in part **a**, excepting that 2 μM UBE2N/UBE2V2 (**b**), UBE2K (**c**), or UBE2D1 (**d**) was used, with recombinant TRIM21 RING (R), TRIM2 RBCC, or TRIM23 RBB as positive controls alongside either TRIM6 or TRIM22 R or RB (n=3). **e**, Western blots of auto-ubiquitination reaction carried out at 30 °C 120 min using 1 μM UBE2D1, 1 μM UBA1, 50 μM ubiquitin, and 3 mM ATP, using FLAG pull downs from either untransfected, FLAG-TRIM2, 6, or 22 transfected HEK293T cells that were either untreated or treated 18 h with 5,000 units/ml IFN-β or 20 ng/ml TNFα (n=3). **f**, Quantification of n=3 independent experiments as described in part **e**, with individual values plotted as circles, error bars represent mean ± SEM. **g**, Volcano plots indicating the IFN-β-responsive interactomes of FLAG-TRIM6 (left) and FLAG-TRIM22 (right), as determined by timsTOF mass spectrometry of anti-FLAG immunoprecipitated complexes from transfected HEK293T cells (5,000 units/ml IFN-β 18 h vs. untreated), with TRIM21 identified as a hit in both cases (teal). **h**, Volcano plot of global proteomic changes in HEK293T cells following treatment with 5,000 units/ml IFN-β 18 h. **i**, Pull down using 'GFP clamp' DARPin pull down against GFP-tagged TRIM6 and TRIM22 expressed in HEK293T cells, with or without pre-treatment with 5,000 units/ml IFN-β 18 h, before use in a ubiquitination reaction as described in part **e**.

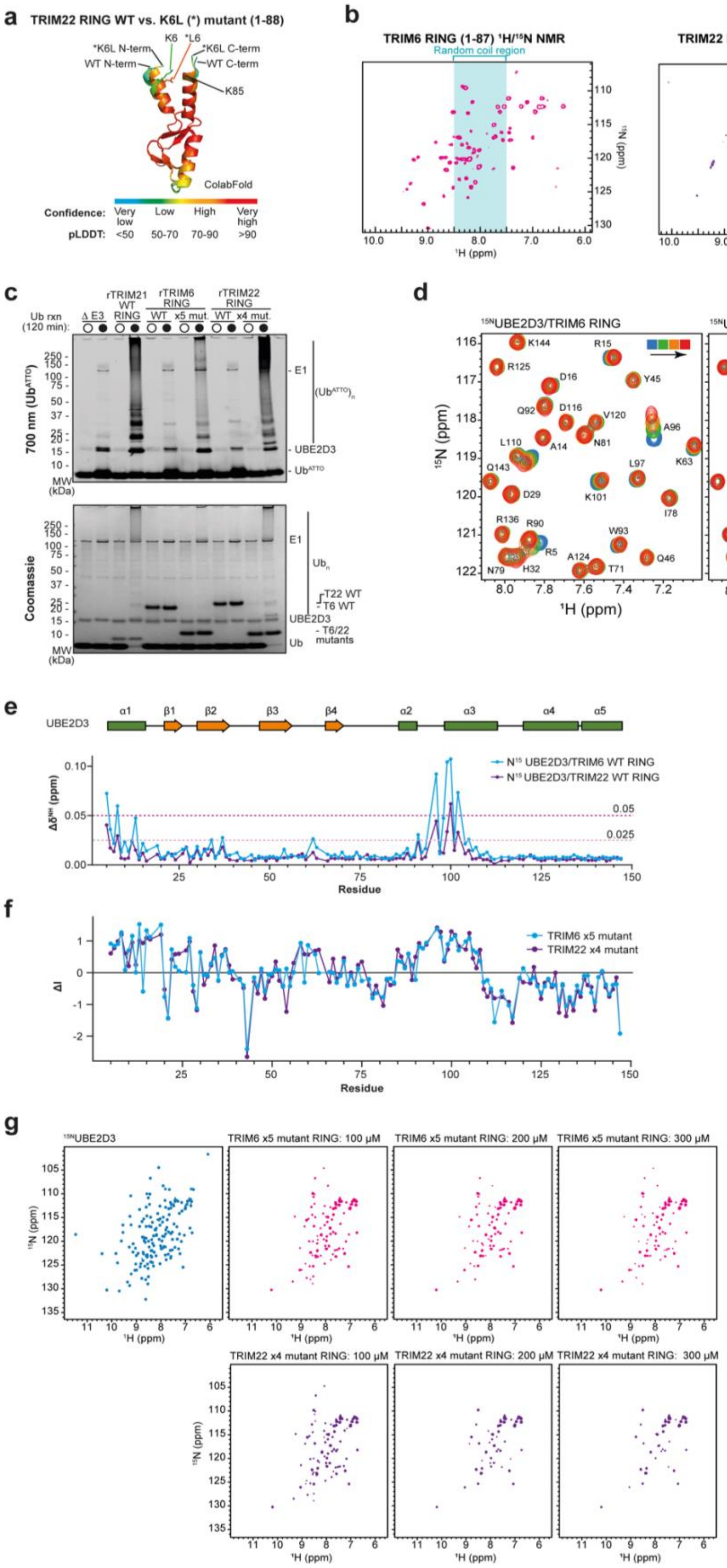
a
TRIM22 RING WT vs. K6L (*) mutant (1-88)
ColabFold
Confidence: Very low Low High Very high
pLDDT: <50 50-70 70-90 >90
b
TRIM6 RING (1-87) ¹H/¹⁵N NMR
TRIM22 RING (1-87) ¹H/¹⁵N NMR
Random coil region
c
d
¹⁵NUBE2D3/TRIM6 RING
¹⁵NUBE2D3/TRIM22 RING
e
f
TRIM6 x5 mutant
TRIM22 x4 mutant
g
TRIM6 x5 mutant RING: 100 µM
TRIM6 x5 mutant RING: 200 µM
TRIM6 x5 mutant RING: 300 µM
TRIM22 x4 mutant RING: 100 µM
TRIM22 x4 mutant RING: 200 µM
TRIM22 x4 mutant RING: 300 µM

**Supplementary Figure 6 – Data supporting key interfaces with E2~ubiquitin are mutated in TRIM6 and TRIM22, leading to lack of ubiquitin ligase activity *in vitro*. a**, Alignments of ColabFold[70] predictions of wild-type (WT) and K6L mutant TRIM22 (residues 1-88), demonstrating increased confidence in well-folded and closer $\alpha$1 and $\alpha$3 helices. **b**, $^{1}$H-$^{15}$N HSQC spectra of TRIM6 (left, pink) and TRIM22 (right, purple) RING domains, with the spectral regions corresponding to resonances characteristic of residues in random coil conformation indicated in teal. **c**, 700 nm fluorescent imaging and Coomassie stains of auto-ubiquitination reactions carried out at 30 °C 120 min using 2 μM UBE2D3 with 1 μM UBA1, 50 μM ubiquitin, 1 μM $Ub^{ATTO}$ and 3 mM ATP, with recombinantly purified full-length $His_6$-SUMO-tagged wild-type (WT) or $His_6$-tagged active mutant (TRIM6 P41A/N42V/G43I/R54S/Q60R 'x5 mutant'; TRIM22 K6L/K42V/Q60R/K85V 'x4 mutant') TRIM6 or TRIM22 (n=3). **d**, Details of the $^{1}$H-$^{15}$N HSQC spectra of UBE2D3 titrated with the RING domain of TRIM6 and TRIM22 WT. Spectra from blue to red at different ligand concentrations (0, 100, 200, 300 μM) are plotted at the same contour level. **e**, Chemical shift perturbations ($\Delta\delta^{NH}$) versus residue number in the NMR spectra of $^{15}$N-labelled UBE2D3 in the presence of two molar equivalents (300 μM) of the RING domains of TRIM6 (blue) and TRIM22 (purple). Secondary structure of UBE2D3 is reported as a function of residue number. **f**, Residue-specific values of differential line broadening ($\Delta$I) of a 150 μM sample of $^{15}$N-labelled UBE2D3 induced by x5 mutant TRIM6 (blue) and x4 mutant TRIM22 (purple) at 100 μM concentration of RING domains. Residues with positive values above $\Delta$I were mapped on the surface of UBE2D3 (Fig. 4h). **g**, Side-by-side comparison of the $^{1}$H-$^{15}$N HSQC spectra of $^{15}$N-labelled UBE2D3 in the absence (blue) or presence of the RING domain of x5 mutant TRIM6 (pink) and x4 mutant TRIM22 (purple) at increasing concentrations. Source data is provided as a Source Data file.

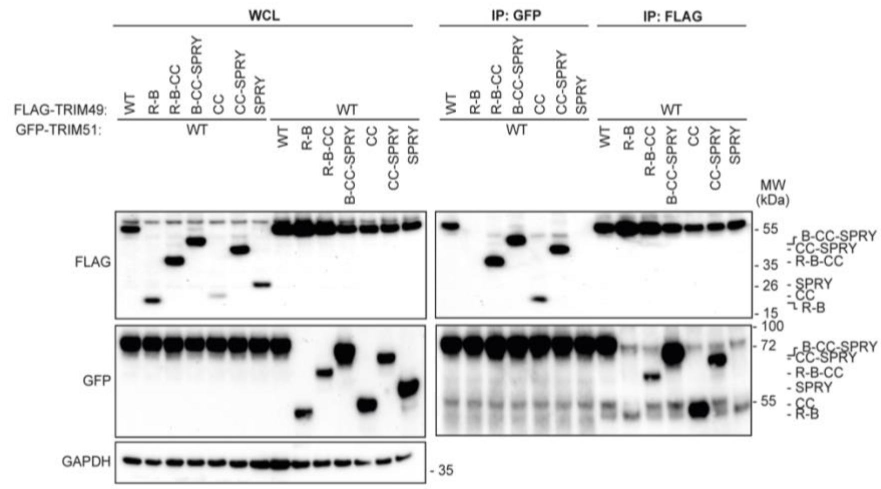

**Supplementary Figure 7 – Data supporting TRIM49 and TRIM51 form an active:inactive pair that cross-regulate, with conflicting effects on autophagy**

Western blot analysing the domains of FLAG-TRIM49 and GFP-TRIM51 that interact using co-immunoprecipitation of the indicated truncation mutant proteins from HEK293T cells (n=3, see **Fig. 5h**).

### Supplementary Tables

**Supplementary Table 1 –** Comparative analysis of TRIM localisation identified in this study compared to that found previously in the literature, with any discrepancies highlighted in grey and the likely cause suggested

| TRIM | Localisation | | |
|---|---|---|---|
| | **Literature** | **This study** | **Cause of discrepancy** (listed respective to literature references) |
| 1 | Cytoskeleton[1-4] | Diffuse, nuclear | Different cell types (HeLa, H1299, Cos7, COS-1 cells) |
| 2 | Cytoskeleton[1,5] | Cytoskeleton | - |
| 3 | Diffuse[1,6] | Diffuse | - |
| 4 | Puncta[1,7], mitochondria[7] | Puncta | - |
| 5 | Puncta[1,8,9] | Puncta | - |
| 6 | Puncta[1,10] | Puncta | - |
| 7 | Diffuse[1,11], puncta[12], nuclear[1] | Puncta | - |
| 8 | Diffuse[13], puncta[1], nuclear[1,13] | Diffuse, nuclear | - |
| 9 | Puncta[1,14] | Puncta, aggregates | - |
| 10 | Diffuse[15], aggregates[1] | Diffuse | - |
| 11 | Diffuse[1], nuclear[1] | Diffuse, nuclear | - |
| 12 | Diffuse[1], puncta[1,16,17] | Puncta | - |
| 13 | Endoplasmic reticulum[1,18] | Puncta | Different cell type (HeLa) or endogenous staining |
| 15 | Puncta[19], cytoskeleton[20] | Puncta, aggregates | - |
| 17 | Diffuse[21] | Diffuse | - |
| 18 | Diffuse[14], cytoskeleton[1,22] | Cytoskeleton | - |
| 19 | Nuclear[1,23] | Nuclear | - |
| 21 | Diffuse[1], puncta[1,24] | Diffuse, puncta | - |
| 22 | Diffuse[1], puncta[1,25], nuclear[25,26] | Nuclear | - |
| 23 | Diffuse[27], puncta[1,27], nuclear[1] | Diffuse | - |
| 24 | Puncta[1], nuclear[28,29] | Nuclear | - |
| 25 | Diffuse[1], puncta[30], aggregates[1] | Puncta | - |
| 26 | Diffuse[1,31,32], puncta[33], nuclear[32] | Diffuse | - |
| 27 | Puncta[1,34], nuclear[34,35] | Puncta | - |
| 28 | Nuclear[1,36,37] | Nuclear | - |
| 30 | Diffuse[17], puncta[1,38], aggregates[17,38] | Diffuse | - |
| 31 | Diffuse[1], puncta[39-41] | Puncta | - |
| 32 | Puncta[1,14,42-45], nuclear[46] | Puncta, aggregates | - |
| 33 | Diffuse[47], nuclear[47-49] | Nuclear | - |
| 34 | Puncta[50] | Puncta | - |
| 35 | Diffuse[51] | Diffuse, nuclear | - |
| 36 | Cytoskeleton[52,53] | Diffuse, nuclear | Different cell type (HeLa) |
| 37 | Diffuse[54-57], puncta[55], aggregates[58], nuclear[56] | Diffuse | - |
| 38 | Puncta[59] | Puncta | - |
| 39 | Diffuse[60] | Diffuse | - |
| 40 | Diffuse[61], nuclear[61] | Diffuse, nuclear | - |
| 41 | Diffuse[62-64], nuclear[62] | Aggregates | Different cell type (SH-SY5Y, A549) with endogenous staining, or different cell type (RAW264.7) |
| 42 | Untested | Puncta | - |
| 43 | Puncta[65] | Diffuse | Different cell type (HFF) stimulated with HSV-1 |
| 45 | Diffuse[66,67] | Diffuse, aggregates | - |
| 46 | Cytoskeleton[52,68,69] | Diffuse, nuclear | Different cell type (HeLa) or endogenous protein in hippocampal neurones |
| 47 | Diffuse[70] | Diffuse, aggregates | - |
| 48 | Untested | Puncta | - |
| 49 | Untested | Diffuse, nuclear | - |
| 50 | Diffuse[71,72], puncta[73-76] | Diffuse | - |

| | | | |
|---|---|---|---|
| 51 | Untested | Diffuse, nuclear | - |
| 52 | Nuclear[77] | Nuclear | - |
| 54 | Untested | Diffuse, aggregates | - |
| 55 | Diffuse[78-80], puncta[78] | Puncta | - |
| 56 | Diffuse[81,82] | Diffuse | - |
| 58 | Diffuse[83,84] | Nuclear | Different cell types (A549, HCC827, or HeLa) and tag (FLAG or mCherry) |
| 59 | Diffuse[85] | Diffuse | - |
| 60 | Aggregates[86] | Diffuse, nuclear | Different cell type (CHO-K1) |
| 61 | Untested | Nuclear | - |
| 62 | Diffuse[87], aggregates[88] | Diffuse, aggregates | - |
| 63 | Puncta[89-91], mitochondria[92] | Puncta | - |
| 64 | Untested | Diffuse, aggregates | - |
| 65 | Diffuse[93-95] | Diffuse, nuclear | - |
| 67 | Diffuse[96,97], puncta[96,97] | Diffuse | - |
| 68 | Diffuse[98] | Diffuse | - |
| 69 | Diffuse[99], aggregates[100,101] | Cytoskeleton | Different cell type (HeLa) and tags (FLAG or Myc) |
| 71 | Puncta[102,103] | Diffuse | Different cell types (KH2, embryocarcinoma, or HeLa cells) with endogenous staining |
| 72 | Diffuse[104,105], nuclear[104] | Diffuse | - |
| 73 | Untested | Diffuse | - |
| 74 | Untested | Diffuse, aggregates | - |
| 75 | Nuclear[106] | Nuclear | - |
| 77 | Untested | Diffuse | - |
| L1 | Untested | Diffuse, nuclear | - |

**Supplementary Table 2 –** Comparative analysis of TRIM ubiquitin ligase activity identified in this study compared to that found previously in the literature, with any discrepancies highlighted in grey and the likely cause suggested

| | *in vitro* auto-ubiquitination activity | | | In cell auto-ubiquitination activity | | |
|---|---|---|---|---|---|---|
| **TRIM** | **Literature** | **This study** | **Likely cause of discrepancy** (listed respective to literature references) | **Literature** | **This study** | **Likely cause of discrepancy** (listed respective to literature references) |
| 1 | Yes[14] | Yes | - | Yes[2,14] | Yes | - |
| 2 | Yes[107] | Yes | - | Yes[107] | No | Different cell type (HeLa) and IP (His-Ub IP) |
| 3 | Yes[108] | No | Lacks key RING dimerisation residues | Yes[6,109,110] | No | Lacks key RING dimerisation residues |
| 4 | Yes[111] | Yes | - | Yes[7] | No | Substrate ubiquitination rather than auto-ubiquitination |
| 5 | Yes[8] | Yes | - | Yes[8,9] | Yes | - |
| 6 | Yes[10] | No | Substrate ubiquitination rather than auto-ubiquitination | Yes[10,112] | Yes | - |
| 7 | Yes[11,113,114] | Yes | - | Yes[11,12,113,114] | No | Substrate ubiquitination rather than auto-ubiquitination |
| 8 | Yes[13] | Yes | - | Yes[13,115] | No | Substrate ubiquitination rather than auto-ubiquitination |
| 9 | Yes[14] | Yes | - | Yes[116] | No | Unknown |
| 10 | Untested | Yes | - | Yes[15,117,118] | No | Substrate ubiquitination rather than auto-ubiquitination |
| 11 | Yes[14] | Yes | - | Yes[14] | Yes | - |
| 12 | Untested | Yes | - | Yes[16] | No | Different isoform (Trim12c rather than Trim12a) |
| 13 | Yes[18] | Yes | - | Yes[18] | Yes | - |
| 15 | Yes[119] | No | Substrate ubiquitination rather than auto-ubiquitination | Yes[120] | No | Substrate ubiquitination rather than auto-ubiquitination |
| 17 | Yes[121] | Yes | - | Yes[21,121] | No | Different cell type (COS7) |
| 18 | Yes[122] | Yes | - | Yes[22] | No | Substrate ubiquitination rather than auto-ubiquitination |
| 19 | Untested | Yes | - | No[23,123] | Yes | Different cell type (HeLa) |
| 21 | Yes[124] | Yes | - | Yes[24,125] | Yes | - |
| 22 | Yes[126] | No | Different E2 (activity shown with Ube2D2) | Yes[26,126] | Yes | - |
| 23 | Yes[127,128] | Yes | - | Yes[27,128-130] | Yes | - |
| 24 | Yes[28] / No[131] | No | - | Yes[28,132,133] | Yes | - |
| 25 | Yes[30,134-136] | Yes | - | Yes[134,135,137] | Yes | - |
| 26 | Yes[138-140] | Yes | - | Yes[31,33,138-141] | Yes | - |
| 27 | Yes[14,35,142,143] | Yes | - | Yes[14,34,142-146] | Yes | - |
| 28 | Yes[147] / No[131] | No | - | Yes[148] | No | Ubiquitin E3 ligase activity under debate |
| 30 | Untested | Yes | - | Yes[149] | Yes | - |
| 31 | Yes[40,41,150-152] | Yes | - | Yes[39,40,150-155] | Yes | - |
| 32 | Yes[14,45,136,156-158] | Yes | - | Yes[14,45,46,156,159,160] | Yes | - |
| 33 | Yes[161] / No[131] | No | - | Yes[161,162] | No | Ubiquitin E3 ligase activity under debate |
| 34 | Yes[163] | Yes | - | Yes[163] | No | Substrate ubiquitination rather than auto-ubiquitination |
| 35 | Yes[164] | Yes | - | Yes[164-167] | Yes | - |
| 36 | Yes[53] | No | Different E2 (activity shown with Ube2D2) | Yes[53,168] | Yes | - |
| 37 | Yes[57,58,169] | No | Different E2 (activity shown with Ube2D2) | Yes[54,55,58,169,170] | Yes | - |
| 38 | Untested | No | - | Yes[171-174] | No | Unknown |
| 39 | Yes[175] | Yes | - | Yes[176-178] | No | Substrate ubiquitination rather than auto-ubiquitination |
| 40 | Yes[179] | No | Substrate ubiquitination rather than auto-ubiquitination | Yes[179-181] | No | Substrate ubiquitination rather than auto-ubiquitination |
| 41 | Yes[63,64] | Yes | - | Yes[62-64,182] | Yes | - |
| 42 | Untested | Yes | - | Untested | Yes | - |

| | | | | | | |
|---|---|---|---|---|---|---|
| 43 | Untested | Yes | - | Yes[65] | No | Substrate ubiquitination rather than auto-ubiquitination |
| 45 | Yes[66,183] | Yes | - | Yes[67,184] | Yes | - |
| 46 | Yes[183] | No | Substrate ubiquitination rather than auto-ubiquitination | Yes[183,185-187] | Yes | - |
| 47 | Yes[188] | No | Different E2 (activity shown with Ube2D2) | Yes[70,188-191] | No | Substrate ubiquitination rather than auto-ubiquitination |
| 48 | Yes[192] | Yes | - | Yes[192] | Yes | - |
| 49 | No[193] | Yes | Mammalian expression untested (purified from bacteria) | Yes[193] | Yes | - |
| 50 | Yes[76] | Yes | - | Yes[71-75,194] | No | Substrate ubiquitination rather than auto-ubiquitination |
| 51 | Untested | No | - | Untested | No | - |
| 52 | Yes[195] | Yes | - | Yes[196-198] | Yes | - |
| 54 | Yes[199,200] | Yes | - | Yes[201,202] | No | Substrate ubiquitination rather than auto-ubiquitination |
| 55 | Yes[203] | Yes | - | Yes[204] | Yes | - |
| 56 | Yes[205,206] | Yes | - | Yes[81,205] | Yes | - |
| 58 | Yes[83] | Yes | - | Yes[83,84,207-210] | Yes | - |
| 59 | Yes[211] | Yes | - | Yes[211-214] | Yes | - |
| 60 | Untested | Yes | - | Untested | Yes | - |
| 61 | Untested | Yes | - | Untested | Yes | - |
| 62 | Yes[88] | Yes | - | Yes[88,215,216] | Yes | - |
| 63 | Yes[90] | Yes | - | Yes[89,90] | Yes | - |
| 64 | Untested | Yes | - | Yes[217] | Yes | - |
| 65 | Yes[93,218] | Yes | - | Yes[93-95,218-220] | Yes | - |
| 67 | Untested | Yes | - | Yes[96,97,221] | No | Substrate ubiquitination rather than auto-ubiquitination |
| 68 | Yes[222] | Yes | - | Yes[98] | No | Substrate ubiquitination rather than auto-ubiquitination |
| 69 | Yes[100,223] | Yes | - | Yes[100,101,224] | No | Unknown |
| 71 | Yes[103] | Yes | - | Yes[103,225,226] | Yes | - |
| 72 | Yes[227,228] | No | Different E2 (activity shown with Ube2D3 or 2C) | Yes[228] | No | Substrate ubiquitination rather than auto-ubiquitination |
| 73 | Untested | Yes | - | Untested | Yes | - |
| 74 | Untested | Yes | - | Untested | Yes | - |
| 75 | Untested | Yes | - | Untested | Yes | - |
| 77 | Untested | Yes | - | Untested | No | - |
| L1 | Untested | Yes | - | Untested | Yes | - |

**Supplementary Table 3 –** TRIM RING sequences and AlphaFold2 references

| TRIM | RING sequence | AlphaFold2 PDB |
|---|---|---|
| TRIM1 | MGESPASVVLNASGGLFSLKMETLESELTCPICLELFEDPLLLPCAHSLCFSCAHRILVSSCSSGESIEPITAFQCPTCRYVISLNHRGLDGLKRNVTLQNIIDRFQKASVSGP | AF-Q9UJV3-F1-model_v4 |
| TRIM2 | MASEGTNIPSPVVRQIDKQFLICSICLERYKNPKVLPCLHTFCERCLQNYIPAHSLTLSCPVCRQTSILPEKGVAALQNNFFITNLMDVLQRTP | AF-Q9C040-F1-model_v4 |
| TRIM3 | MAKREDSPGPEVQPMDKQFLVCSICLDRYQCPKVLPCLHTFCERCLQNYIPAQSLTLSCPVCRQTSILPEQGVSALQNNFFISSLMEAMQQAP | AF-O75382-F1-model_v4 |
| TRIM4 | MEAEDIQEELTCPICLDYFQDPVSIECGHNFCRGCLHRNWAPGGGPFPCPECRHPSAPAALRPNWALARLTEKTQRRRLGPVP | AF-Q9C037-F1-model_v4 |
| TRIM5 | MASGILVNVKEEVTCPICLELLTQPLSLDCGHSFCQACLTANHKKSMLDKGESSCPVCRISYQPENIRPNRHVANIVEKLREVKLSP | AF-Q9C035-F1-model_v4 |
| TRIM6 | MTSPVLVDIREEVTCPICLELLTEPLSIDCGHSFCQACITPNGRESVIGQEGERSCPVCQTSYQPGNLRPNRHLANIVRRLREVVLGP | AF-Q9C030-F1-model_v4 |
| TRIM7 | MAAVGPRTGPGTGAEALALAAELQGEATCSICLELFREPVSVECGHSFCRACIGRCWERPGAGSVGAATRAPPFPLPCPQCREPARPSQLRPNRQLAAVATLLRRFSLP | AF-Q9C029-F1-model_v4 |
| TRIM8 | MAENWKNCFEEELICPICLHVFVEPVQLPCKHNFCRGCIGEAWAKDSGLVRCPECNQAYNQKPGLEKNLKLTNIVEKFNALHVEKP | AF-Q9BZR9-F1-model_v4 |
| TRIM9 | MEEMEEELKCPVCGSFYREPIILPCSHNLCQACARNILVQTPESESPQSHRAAGSGVSDYDYLDLDKMSLYSEADSGYGSYGGFASAPTTPCQKSPNGVRVFPPAMPPPATHLSPALAPVPRNSCITCPQCHRSLILDDRGLRGFPKNRVLEGVIDRYQQSKA | AF-Q9C026-F1-model_v4 |
| TRIM10 | MASAASVTSLADEVNCPICQGTLREPVTIDCGHNFCRACLTRYCEIPGPDLEESPTCPLCKEPFRPGSFRPNWQLANVVENIERLQLVSTLGLGE | AF-Q9UDY6-F1-model_v4 |
| TRIM11 | MAAPDLSTNLQEEATCAICLDYFTDPVMTDCGHNFCRECIRRCWGQPEGPYACPECRELSPQRNLRPNRPLAKMAEMARRLHP | AF-Q96F44-F1-model_v4 |
| Trim12 | MASQFMKNLKEEVTCPVCLNLMVKPVSADCGHTFCQGCITLYFESIKCDKKVFICPVCRISYQFSNLRPNRNVANIVERLKMFKPSP | AF-Q99PQ1-F1-model_v4 |
| TRIM13 | MELLEEDLTCPICCSLFDDPRVLPCSHNFCKKCLEGILEGSVRNSLWRPAPFKCPTCRKETSATGINSLQVNYSLKGIVEKYNKIKISP | AF-O60858-F1-model_v4 |
| TRIM15 | MPATPSLKVVHELPACTLCAGPLEDAVTIPCGHTFCRLCLPALSQMGAQSSGKILLCPLCQEEEQAETPMAPVPLGPLGETYCE | AF-Q9C019-F1-model_v4 |
| TRIM17 | MEAVELARKLQEEATCSICLDYFTDPVMTTCGHNFCRACIQLSWEKARGKKGRRKRKGSFPCPECREMSPQRNLLPNRLLTKVAEMAQQHP | AF-Q9Y577-F1-model_v4 |
| TRIM18 | METLESELTCPICLELFEDPLLLPCAHSLCFNCAHRILVSHCATNESVESITAFQCPTCRHVITLSQRGLDGLKRNVTLQNIIDRFQKASVSGP | AF-O15344-F1-model_v4 |
| TRIM19 | MEPAPARSPRPQQDPARPQEPTMPPPETPSEGRQPSPSPSPTERAPASEEEFQFLRCQQCQAEAKCPKLLPCLHTLCSGCLEASGMQCPICQAPWPLGADTPALDNVFFESLQRRLSVYRQIVDAQ | AF-P29590-F1-model_v4 |
| TRIM21 | MASAARLTMMWEEVTCPICLDPFVEPVSIECGHSFCQECISQVGKGGGSVCPVCRQRFLLKNLRPNRQLANMVNNLKEISQEARE | AF-P19474-F1-model_v4 |
| TRIM22 | MDFSVKVDIEKEVTCPICLELLTEPLSLDCGHSFCQACITAKIKESVIISRGESSCPVCQTRFQPGNLRPNRHLANIVERVKEVKMSP | AF-Q8IYM9-F1-model_v4 |
| TRIM23 | MATLVVNKLGAGVDSGRQGSRGTAVVKVLECGVCEDVFSLQGDKVPRLLLCGHTVCHDCLTRLPLHGRAIRCPFDRQVTDLGDSGVWGLKKNFALLELLERLQNGP | AF-P36406-F1-model_v4 |
| TRIM24 | MEVAVEKAVAAAAAASAAASGGPSAAPSGENEAESRQGPDSERGGEAARLNLLDTCAVCHQNIQSRAPKLLPCLHSFCQRCLPAPQRYLMLPAPMLGSAETPPPVPAPGSPVSGSSPFATQVGVIRCPVCSQECAERHIIDNFFVKDTTEVP | AF-O15164-F1-model_v4 |
| TRIM25 | MAELCPLAEELSCSICLEPFKEPVTTPCGHNFCGSCLNETWAVQGSPYLCPQCRAVYQARPQLHKNTVLCNVVEQFLQADLAREP | AF-Q14258-F1-model_v4 |
| TRIM26 | MATSAPLRSLEEEVTCSICLDYLRDPVTIDCGHVFCRSCTTDVRPISGSRPVCPLCKKPFKKENIRPVWQLASLVENIERLKVDKGRQP | AF-Q12899-F1-model_v4 |
| TRIM27 | MASGSVAECLQQETTCPVCLQYFAEPMMLDCGHNICCACLARCWGTAETNVSCPQCRETFPQRHMRPNRHLANVTQLVKQLRTERP | AF-P14373-F1-model_v4 |
| TRIM28 | MAASAAAASAAAASAASGSPGPGEGSAGGEKRSTAPSAAASASASAAASSPAGGGAEALELLEHCGVCRERLRPEREPRLLPCLHSACSACLGPAAPAAANSSGDGGAAGDGTVVDCPVCKQQCFSKDIVENYFMRDSGSKAATDA | AF-Q13263-F1-model_v4 |
| Trim30 | MASSVLEMIKEEVTCPICLELLKEPVSADCNHSFCRACITLNYESNRNTDGKGNCPVCRVPYPFGNLRPNLHVANIVERLKGFKSIP | AF-P15533-F1-model_v4 |
| TRIM31 | MASGQFVNKLQEEVICPICLDILQKPVTIDCGHNFCLKCITQIGETSCGFFKCPLCKTSVRKNAIRFNSLLRNLVEKIQALQASEVQSKRKE | AF-Q9BZY9-F1-model_v4 |
| TRIM32 | MAAAAASHLNLDALREVLECPICMESFTEEQLRPKLLHCGHTICRQCLEKLLASSINGVRCPFCSKITRITSLTQLTDNLTVLKIIDTAGLSEAVGLLMCRSCGRRLP | AF-Q13049-F1-model_v4 |
| TRIM33 | MAENKGGGEAESGGGGSGSAPVTAGAAGPAAQEAEPPLTAVLVEEEEEEGGRAGAEGGAAGPDDGGVAAASSGSAQAASSPAASVGTGVAGGAVSTPAPAPASAPAPGPSAGPPPGPPASLLDTCAVCQQSLQSRREAEPKLLPCLHSFCLRCLPEPERQLSVPIPGGSNGDIQQVGVIRCPVCRQECRQIDLVDNYFVKDTSEAP | AF-Q9UPN9-F1-model_v4 |
| TRIM34 | MASKILLNVQEEVTCPICLELLTEPLSLDCGHSLCRACITVSNKEAVTSMGGKSSCPVCGISYSFEHLQANQHLANIVERLKEVKLSP | AF-Q9BYJ4-F1-model_v4 |
| TRIM35 | MERSPDVSPGPSRSFKEELLCAVCYDPFRDAVTLRCGHNFCRGCVSRCWEVQVSPTCPVCKDRASPADLRTNHTLNNLVEKLLREEAEGARWTSYR | AF-Q9UPQ4-F1-model_v4 |
| TRIM36 | MSESGEMSEFGYIMELIAKGKVTIKNIERELICPACKELFTHPLILPCQHSICHKCVKELLLTLDDSFNDVGSDNSNQSSPRLRLPSPSMDKIDRINRPGWKRNSLTPRTTVFPCPGCEHDVDLGERGINGLFRNFTLETIVERYRQAARA | AF-Q9NQ86-F1-model_v4 |
| TRIM37 | MDEQSVESIAEVFRCFICMEKLRDARLCPHCSKLCCFSCIRRWLTEQRAQCPHCRAPLQLRELVNCRWAEEVTQQLDTLQLCSLTKHE | AF-O94972-F1-model_v4 |
| TRIM38 | MASTTSTKKMMEEATCSICLSLMTNPVSINCGHSYCHLCITDFFKNPSQKQLRQETFCCPQCRAPFHMDSLRPNKQLGSLIEALKETDQE | AF-O00635-F1-model_v4 |
| TRIM39 | MAETSLLEAGASAASTAAALENLQVEASCSVCLEYLKEPVIIECGHNFCKACITRWWEDLERDFPCPVCRKTSRYRSLRPNRQLGSMVEIAKQLQAVKRKIRDE | AF-Q9HCM9-F1-model_v4 |
| TRIM40 | MIPLQKDNQEEGVCPICQESLKEAVSTNCGHLFCRVCLTQHVEKASASGVFCCPLCRKPCSEEVLG | AF-Q6P9F5-F1-model_v4 |
| TRIM41 | MAAVAMTPNPVQTLQEEAVCAICLDYFTDPVSIGCGHNFCRVCVTQLWGGEDEEDRDELDREEEEEDGEEEEVEAVGAGAGWDTPMRDEDYEGDMEEEVEEEEEGVFWTSGMSRSSWDNMDYVWEEEDEEEDLDYYLGDMEEEDLRGEDEEDEEEVLEEVEEEDLDPVTPLPPPPAPRRCFTCPQCRKSFPRRSFRPNLQLANMVQVIRQMHP | AF-Q8WV44-F1-model_v4 |
| TRIM42 | METAMCVCCPCCTWQRCCPQLCSCLCCKFIFTSERNCTCFPCPYKDERNCQFCHCTCSESPNCHWCCCSWANDPNCKCCCTASSNLNCYYYESRCCRNTIITFHKGRLRSIHTSSKTALRTGSSDTQVDEVKSIPANSHLVNHLNCPMCSRLRLHSFMLPCNHSLCEKCLRQLQKHAEVTENFFILICPVCDRSHCMPYSNKMQLPENYLHGRLTKRYMQEHGYLKWRFDRSSGP | AF-Q8IWZ5-F1-model_v4 |

| | | |
|---|---|---|
| TRIM43 | MDSDFSHAFQKELTCVICLNYLVDPVTICCGHSFCRPCLCLSWEEAQSPANCP<br>ACREPSPKMDFKTNILLKNLVTIARKASLWQFLSSE | AF-Q96BQ3-F1-model_v4 |
| TRIM45 | MSENRKPLLGFVSKLTSGTALGNSGKTHCPLCLGLFKAPRLLPCLHTVCTTCLE<br>QLEPFSVVDIRGGDSDTSSEGSIFQELKPRSLQSQIGILCPVCDAQVDLPMGGV<br>KALTIDHLAVNDVMLESLRGE | AF-Q9H8W5-F1-model_v4 |
| TRIM46 | MAEGEDMQTFTSIMDALVRISTSMKNMEKELLCPVCQEMYKQPLVLPCTHNVC<br>QACAREVLGQQGYIGHGGDPSSEPTSPASTPSTRSPRLSRRTLPKPDRLDRLL<br>KSGFGTYPGRKRGALHPQVIMFPCPACQGDVELGERGLAGLFRNLTLERVVER<br>YRQSVSVG | AF-Q7Z4K8-F1-model_v4 |
| TRIM47 | MDGSGPFSCPICLEPLREPVTLPCGHNFCLACLGALWPHRGASGAGGPGGAA<br>RCPLCQEPFPDGLQLRKNHTLSELLQLRQGSGP | AF-Q96LD4-F1-model_v4 |
| TRIM48 | MSRRIIVGTLQRTQRNMNSGISQVFQRELTCPICMNYFIDPVTIDCGHSFCRPCF<br>YLNWQDIPILTQCFECIKTIQQRNLKTNIRLKKMASLARKASLWLFLSSE | AF-Q8IWZ4-F1-model_v4 |
| TRIM49 | MNSGILQVFQGELICPLCMNYFIDPVTIDCGHSFCRPCFYLNWQDIPFLVQCSE<br>CTKSTEQINLKTNIHLKKMASLARKVSLWLFLSSE | AF-P0CI25-F1-model_v4 |
| TRIM50 | MAWQVSLPELEDRLQCPICLEVFKEPLMLQCGHSYCKGCLVSLSCHLDAELRC<br>PVCRQAVDGSSSLPNVSLARVIEALRLP | AF-Q86XT4-F1-model_v4 |
| TRIM51 | MNSGILQVFQRALTCPICMNYFLDPVTIDCGHSFCRPCLYLNWQDTAVLAQCSE<br>CKKTTRQRNLNTDICLKNMAFIARKASLRQFLSSE | AF-Q9BSJ1-F1-model_v4 |
| TRIM52 | MAGYATTPSPMQTLQEEAVCAICLDYFKDPVSISCGHNFCRGCVTQLWSKEDE<br>EDQNEEEDEWEEEEDEEAVGAMDGWDGSIREVLYRGNADEELFQDQDDDEL<br>WLGDSGITNWDNVDYMWDEEEEEEEEDQDYYLGGLRPDLRIDVYREEEILEAY<br>DEDEDEELYPDIHPPPSLPLPGQFTCPQCRKSFTRRSFRPNLQLANMVQIIRQM<br>CP | AF-Q96A61-F1-model_v4 |
| TRIM54 | MNFTVGFKPLLGDAHSMDNLEKQLICPICLEMFSKPVVILPCQHNLCRKCANDV<br>FQASNPLWQSRGSTTVSSGGRFRCPSCRHEVVLDRHGVYGLQRNLLVENIIDI<br>YKQESSRP | AF-Q9BYV2-F1-model_v4 |
| TRIM55 | MSASLNYKSFSKEQQTMDNLEKQLICPICLEMFTKPVVILPCQHNLCRKCASDIF<br>QASNPYLPTRGGTTMASGGRFRCPSCRHEVVLDRHGVYGLQRNLLVENIIDIYK<br>QESTRP | AF-Q9BYV6-F1-model_v4 |
| TRIM56 | MVSHGSSPSLLEALSSDFLACKICLEQLRAPKTLPCLHTYCQDCLAQLADGGRV<br>RCPECRETVPVPPEGVASFKTNFFVNGLLDLVKARACGDLRAG | AF-Q9BRZ2-F1-model_v4 |
| TRIM58 | MAWAPPGERLREDARCPVCLDFLQEPVSVDCGHSFCLRCISEFCEKSDGAQG<br>GVYACPQCRGPFRPSGFRPNRQLAGLVESVRRLGLG | AF-Q8NG06-F1-model_v4 |
| TRIM59 | MHNFEEELTCPICYSIFEDPRVLPCSHTFCRNCLENILQASGNFYIWRPLRIPLK<br>CPNCRSITEIAPTGIESLPVNFALRAIIEKYQQEDHP | AF-Q8IWR1-F1-model_v4 |
| TRIM60 | MEFVTALVNLQEESSCPICLEYLKDPVTINCGHNFCRSCLSVSWKDLDDTFPCP<br>VCRFCFPYKSFRRNPQLRNLTEIAKQLQIRRSKRKRQKE | AF-Q495X7-F1-model_v4 |
| TRIM61 | MEFVTALADLRAEASCPICLDYLKDPVTISCGHNFCLSCIIMSWKDLHDSFPCPF<br>CHFCCPERKFISNPQLGSLTEIAKQLQIRSKKRKRQE | AF-Q5EBN2-F1-model_v4 |
| TRIM62 | MACSLKDELLCSICLSIYQDPVSLGCEHYFCRRCITEHWVRQEAQGARDCPEC<br>RRTFAEPALAPSLKLANIVERYSSFP | AF-Q9BVG3-F1-model_v4 |
| TRIM63 | MDYKSSLIQDGNPMENLEKQLICPICLEMFTKPVVILPCQHNLCRKCANDIFQAA<br>NPYWTSRGSSVSMSGGRFRCPTCRHEVIMDRHGVYGLQRNLLVENIIDIYKQE<br>CSSRP | AF-Q969Q1-F1-model_v4 |
| TRIM64 | MDSDDLQVFQNELICCICVNYFIDPVTIDCGHSFCRPCLCLCSEEGRAPMRCPS<br>CRKISEKPNFNTNVVLKKLSSLARQTRP | AF-A6NGJ6-F1-model_v4 |
| TRIM65 | MAAQLLEEKLTCAICLGLYQDPVTLPCGHNFCGACIRDWWDRCGKACPECRE<br>PFPDGAELRRNVALSGVLEVVRAGP | AF-Q6PJ69-F1-model_v4 |
| TRIM67 | MEEELKCPVCGSLFREPIILPCSHNVCLPCARTIAVQTPDGEQHLPQPLLLSRGS<br>GLQAGAAAAASLEHDAAAGPACGGAGGSAAGGLGGGAGGGGDHADKLSLYS<br>ETDSGYGSYTPSLKSPNGVRVLPMVPAPPGSSAAAARGAACSSLSSSSSSITC<br>PQCHRSASLDHRGLRGFQRNRLLEAIVQRYQQGRGAVP | AF-Q6ZTA4-F1-model_v4 |
| TRIM68 | MDPTALVEAIVEEVACPICMTFLREPMSIDCGHSFCHSCLSGLWEIPGESQNW<br>GYTCPLCRAPVQPRNLRPNWQLANVVEKVRLLRLHP | AF-Q6AZZ1-F1-model_v4 |
| TRIM69 | MEVSTNPSSNIDPGDYVEMNDSITHLPSKVVIQDITMELHCPLCNDWFRDPLML<br>SCGHNFCEACIQDFWRLQAKETFCPECKMLCQYNNCTFNPVLDKLVEKIKKLP | AF-Q86WT6-F1-model_v4 |
| TRIM71 | MASFPETDFQICLLCKEMCGSPAPLSSNSSASSSSSQTSTSSGGGGGGPGAA<br>ARRLHVLPCLHAFCRPCLEAHRLPAAGGGAAGEPLKLRCPVCDQKVVLAEAAG<br>MDALPSSAFLLSNLLDAVVATADEP | AF-Q2Q1W2-F1-model_v4 |
| TRIM72 | MSAAPGLLHQELSCPLCLQLFDAPVTAECGHSFCRACLGRVAGEPAADGTVLC<br>PCCQAPTRPQALSTNLQLARLVEGLAQVP | AF-Q6ZMU5-F1-model_v4 |
| TRIM73 | MAWQVSLLELEDRLQCPICLEVFKESLMLQCGHSYCKGCLVSLSYHLDTKVRC<br>PMCWQVVDGSSSLPNVSLAWVIEALRLP | AF-Q86UV7-F1-model_v4 |
| TRIM74 | MAWQVSLLELEDWLQCPICLEVFKESLMLQCGHSYCKGCLVSLSYHLDTKVRC<br>PMCWQVVDGSSSLPNVSLAWVIEALRLP | AF-Q86UV6-F1-model_v4 |
| TRIM75 | MAVAAALTGLQAEAKCSICLDYLSDPVTIECGHNFCRSCIQQSWLDLQELFPCP<br>VCRHQCQEGHFRSNTQLGRMIEIAKLLQSTKSNKRKQE | AF-A6NK02-F1-model_v4 |
| TRIM77 | MASAITQCSTSELTCSICTDYLTDPVTICCGHRFCSPCLCLLWEDTLTPNCCPVC<br>REISQQMYFKRIIFAEKQVIPTRESVP | AF-I1YAP6-F1-model_v4 |
| TRIML1 | MSTADLMENLREELTCFICLDYFSSPVTTECGHSFCLVCLLRSWEEHNTPLSCP<br>ECWRTLEGPHFQSNERLGRLASIARQLRSQVLQSEDEQGSYGRMP | AF-Q8N9V2-F1-model_v4 |

**Supplementary Table 4 –** Plasmid and antibody reagents used in this study

| **Antibody Database** | | | | |
|---|---|---|---|---|
| **Antibody** | **Use** | **Manufacturer** | **Catalogue Ref** | **Clone** |
| αGFP | Western, IP | Roche | 11814460001 | 7.1 and 13.1 |
| αUbiquitin (FK2: conjugated ubiquitin-specific) | Western | Sigma | ST1200 | FK2 |
| αUbiquitin-HRP (FK2: conjugated ubiquitin-specific) | ELISA | Generon | SMC-214D-HRP | FK2 |
| αFLAG-HRP | Western | Merck | A8592 | Clone M2 |
| αFLAG | IF and IP | Merck | F1804 | Clone M2 |
| αUbiquitin (Ubi: pan-ubiquitin) | Western | Invitrogen | 13-1600 | Ubi-1 |
| αTRIM6 | Western, IF | Protein Tech | 11953-1-AP | Polyclonal |
| αTRIM22 | Western, IF | Atlas Antibodies | HPA003575 | Polyclonal |
| αGAPDH | Western | Millipore | MAB374 | Clone 6C5 |
| αHA-HRP | Western | Merck | 12013819001 | Clone 3F10 |
| αLC3B | Western, IF | Sigma | L7543 | Polyclonal |
| αRabbit-HRP | Western | Cell Signaling | 7074 | Polyclonal |
| αMouse-HRP | Western | Cell Signaling | 7076 | Polyclonal |
| αMouse-HRP | Western | Dako | P0447 | Polyclonal |
| αMouse-Alexa594 | IF | ThermoFisher | A11032 | Polyclonal |
| αRabbit-Alexa488 | IF | ThermoFisher | A11008 | Polyclonal |
| **Plasmid Database** | | | | |
| **Plasmid** | **NCBI Protein Isoform Ref.** | **~Mol. Weight** (untagged kDa) | **Species** | **Derivation (and source)** |
| **Bacterial expression** | | | | |
| pET28-His$_6$-UBA1 | NP_003325.2 | 118 | *Homo sapiens* | N/A (Rittinger Lab, Francis Crick Institute[229]) |
| pET52-His$_6$-UBE2C | NP_008950.1 | 20 | *Homo sapiens* | N/A (Rittinger Lab, Francis Crick Institute[229]) |
| pGEX-6P1-GST-UBE2D1 | NP_003329.1 | 17 | *Homo sapiens* | N/A (Rittinger Lab, Francis Crick Institute[229]) |
| pET49b-GST-UBE2D2 | NP_862821.1 | 17 | *Homo sapiens* | N/A (Rittinger Lab, Francis Crick Institute[131]) |
| pET49b-His$_6$-UBE2D3 | NP_003331.1 | 17 | *Homo sapiens* | N/A (Rittinger Lab, Francis Crick Institute[136]) |
| pET49b-His$_6$-UBE2E1 | NP_003332.1 | 21 | *Homo sapiens* | N/A (Rittinger Lab, Francis Crick Institute[131]) |
| pET49b-GST-UBE2G2 | NP_003334.2 | 19 | *Homo sapiens* | N/A (Rittinger Lab, Francis Crick Institute[229]) |
| pET49b-His$_6$-UBE2K | NP_005330.1 | 22 | *Homo sapiens* | N/A (Rittinger Lab, Francis Crick Institute[131]) |
| pNH-CTH-His$_6$-UBE2N | NP_003339.1 | 17 | *Homo sapiens* | N/A (Rittinger Lab, Francis Crick Institute[229]) |
| pGEX-6P1-GST-UBE2V2 | NP_003341.1 | 16 | *Homo sapiens* | N/A (Rittinger Lab, Francis Crick Institute[229]) |
| pET22b-His$_6$-UBE2W | NP_001001481.3 | 17 | *Homo sapiens* | N/A (Rittinger Lab, Francis Crick Institute[229]) |
| pET49b-His$_6$-TRIM6 RING | NP_477514.1 | 10 | *Homo sapiens* | N/A (Rittinger Lab, Francis Crick Institute[229]) |
| pET49b-His$_6$-TRIM6 RING P41A/N42V/G43I/R54S/Q60R | N/A (mutant) | 10 | N/A (mutant) | This study (see Methods) |
| pET49b-His$_6$-TRIM6 RB | NP_477514.1 | 15 | *Homo sapiens* | N/A (Rittinger Lab, Francis Crick Institute[229]) |
| pET49b-His$_6$-TRIM6 BCC | NP_477514.1 | 12 | *Homo sapiens* | N/A (Rittinger Lab, Francis Crick Institute[229]) |
| pET49b-His$_6$-Trx-TRIM6 RING | NP_477514.1 | 10 | *Homo sapiens* | N/A (Rittinger Lab, Francis Crick Institute[229]) |
| pET49b-His$_6$-TRIM22 RING | NP_006065.2 | 10 | *Homo sapiens* | N/A (Rittinger Lab, Francis Crick Institute[229]) |
| pET49b-His$_6$-TRIM22 RING K6L/K42V/Q60R/K85V | N/A (mutant) | 10 | N/A (mutant) | This study (see Methods) |
| pET49b-His$_6$-TRIM22 RB | NP_006065.2 | 15 | *Homo sapiens* | N/A (Rittinger Lab, Francis Crick Institute[229]) |
| pET49b-His$_6$-TRIM22 BCC | NP_006065.2 | 13 | *Homo sapiens* | N/A (Rittinger Lab, Francis Crick Institute[229]) |
| pET49b-His$_6$-SUMO-TRIM22 RING | NP_477514.1 | 10 | *Homo sapiens* | N/A (Rittinger Lab, Francis Crick Institute[229]) |
| pET49b-His$_6$-TRIM21 RING | NP_003132.2 | 10 | *Homo sapiens* | N/A (Rittinger Lab, Francis Crick Institute[229]) |

| | | | | |
|---|---|---|---|---|
| pET49b-His$_6$-TRIM23 RBB | NP_001647.1 | 23 | *Homo sapiens* | N/A (Rittinger Lab, Francis Crick Institute[229]) |
| pET49b-His$_6$-TRIM15 R | NP_150232.2 | 7 | *Homo sapiens* | This study (see Methods) |
| **Mammalian expression** | | | | |
| pCMV-HA-ubiquitin | NP_001029102.1 | 9 | *Homo sapiens* | N/A (Dundee PPU, #DU48359) |
| pcDNA3.1-FLAG-TRIM15 | NP_150232.2 | 52 | *Homo sapiens* | Sub-cloned from ptCMV-EGFP-TRIM15 (see below) |
| pcDNA3.1-FLAG-TRIM49 | NP_065091.1 | 53 | *Homo sapiens* | Sub-cloned from ptCMV-EGFP-TRIM49 (see below) |
| pcDNA3.1-FLAG-TRIM51 | NP_116070.2 | 53 | *Homo sapiens* | Sub-cloned from ptCMV-EGFP-TRIM49 (see below) |
| pcDNA3.1-FLAG-TRIM49 RB (1-128) | NP_065091.1 | 15 | *Homo sapiens* | Sub-cloned from ptCMV-EGFP-TRIM49 (see below) |
| pcDNA3.1-FLAG-TRIM49 RBCC (1-259) | NP_065091.1 | 31 | *Homo sapiens* | Sub-cloned from ptCMV-EGFP-TRIM49 (see below) |
| pcDNA3.1-FLAG-TRIM49 BCCSPRY (88-450) | NP_065091.1 | 43 | *Homo sapiens* | Sub-cloned from ptCMV-EGFP-TRIM49 (see below) |
| pcDNA3.1-FLAG-TRIM49 CC (129-259) | NP_065091.1 | 16 | *Homo sapiens* | Sub-cloned from ptCMV-EGFP-TRIM49 (see below) |
| pcDNA3.1-FLAG-TRIM49 CCSPRY (129-450) | NP_065091.1 | 37 | *Homo sapiens* | Sub-cloned from ptCMV-EGFP-TRIM49 (see below) |
| pcDNA3.1-FLAG-TRIM49 SPRY (260-450) | NP_065091.1 | 22 | *Homo sapiens* | Sub-cloned from ptCMV-EGFP-TRIM49 (see below) |
| ptCMV-EGFP-TRIM51 RB (1-128) | NP_116070.2 | 15 | *Homo sapiens* | Sub-cloned from ptCMV-EGFP-TRIM51 (see below) |
| ptCMV-EGFP-TRIM51 RBCC (1-259) | NP_116070.2 | 31 | *Homo sapiens* | Sub-cloned from ptCMV-EGFP-TRIM51 (see below) |
| ptCMV-EGFP-TRIM51 BCCSPRY (88-450) | NP_116070.2 | 43 | *Homo sapiens* | Sub-cloned from ptCMV-EGFP-TRIM51 (see below) |
| ptCMV-EGFP-TRIM51 CC (129-259) | NP_116070.2 | 16 | *Homo sapiens* | Sub-cloned from ptCMV-EGFP-TRIM51 (see below) |
| ptCMV-EGFP-TRIM51 CCSPRY (129-450) | NP_116070.2 | 37 | *Homo sapiens* | Sub-cloned from ptCMV-EGFP-TRIM51 (see below) |
| ptCMV-EGFP-TRIM51 SPRY (260-450) | NP_116070.2 | 22 | *Homo sapiens* | Sub-cloned from ptCMV-EGFP-TRIM51 (see below) |
| pcDNA3.1-FLAG-TRIM49 β3-β4 TRIM51 swap | N/A (mutant) | 53 | *Homo sapiens* | gBlock (IDT) of TRIM51 β3-β4 inserted by Gibson Assembly into WT TRIM49 vector, opened by PCR primers flanking the relevant region |
| pcDNA3.1-FLAG-TRIM51 β3-β4 TRIM49 swap | N/A (mutant) | 53 | *Homo sapiens* | gBlock (IDT) of TRIM49 β3-β4 inserted by Gibson Assembly into WT TRIM51 vector, opened by PCR primers flanking the relevant region |
| ptCMV-EGFP-empty | N/A | N/A | N/A | N/A (T. Thurston Laboratory, Imperial College London[230]) |
| ptCMV-EGFP-TRIM1 | NP_036348.2 | 55 | *Homo sapiens* | PCR from 293ET human cDNA, ptCMV-EGFP plasmid derived from pEGFP-N1 plasmid (Clontech) (T. Thurston Laboratory, Imperial College London[230]) |
| ptCMV-EGFP-TRIM2 | NP_001123539.1 | 82 | *Homo sapiens* | PCR from 293ET human cDNA , ptCMV-EGFP plasmid derived from pEGFP-N1 plasmid (Clontech) (T. Thurston Laboratory, Imperial College London[230]) |
| ptCMV-EGFP-TRIM3 | NP_001234935.1 | 82 | *Homo sapiens* | Subcloned from pcDNA3.1-FLAG-TRIM3 (Rittinger Lab, Francis Crick Institute[229]), ptCMV-EGFP plasmid derived from pEGFP-N1 plasmid (Clontech) (T. Thurston |

| | | | | |
|---|---|---|---|---|
| | | | | Laboratory, Imperial College London[230]) |
| ptCMV-EGFP-TRIM4 | NP_148977.2 | 57 | *Homo sapiens* | PCR from mixed human cDNA, ptCMV-EGFP plasmid derived from pEGFP-N1 plasmid (Clontech) (T. Thurston Laboratory, Imperial College London[230]) |
| ptCMV-EGFP-TRIM5 | NP_149023.2 | 55 | *Homo sapiens* | PCR from mixed human cDNA, ptCMV-EGFP plasmid derived from pEGFP-N1 plasmid (Clontech) (T. Thurston Laboratory, Imperial College London[230]) |
| ptCMV-EGFP-TRIM6 | NP_001003818.1 | 56 | *Homo sapiens* | PCR from mixed human cDNA, ptCMV-EGFP plasmid derived from pEGFP-N1 plasmid (Clontech) (T. Thurston Laboratory, Imperial College London[230]) |
| ptCMV-EGFP-TRIM7 | NP_976038.1 | 57 | *Homo sapiens* | PCR from mixed human cDNA, ptCMV-EGFP plasmid derived from pEGFP-N1 plasmid (Clontech) (T. Thurston Laboratory, Imperial College London[230]) |
| ptCMV-EGFP-TRIM8 | NP_112174.2 | 62 | *Homo sapiens* | PCR from mixed human cDNA, ptCMV-EGFP plasmid derived from pEGFP-N1 plasmid (Clontech) (T. Thurston Laboratory, Imperial College London[230]) |
| ptCMV-EGFP-TRIM9 | NP_055978.4 | 80 | *Homo sapiens* | PCR from 293ET human cDNA, ptCMV-EGFP plasmid derived from pEGFP-N1 plasmid (Clontech) (T. Thurston Laboratory, Imperial College London[230]) |
| ptCMV-EGFP-TRIM10 | NP_006769.2 | 55 | *Homo sapiens* | Subcloned from M6P-EGFP vector generated by PCR of mixed human cDNA, ptCMV-EGFP plasmid derived from pEGFP-N1 plasmid (Clontech) (T. Thurston Laboratory, Imperial College London[230]) |
| ptCMV-EGFP-TRIM11 | NP_660215.1 | 53 | *Homo sapiens* | Subcloned from M6P-EGFP vector generated by PCR of mixed human cDNA, ptCMV-EGFP plasmid derived from pEGFP-N1 plasmid (Clontech) (T. Thurston Laboratory, Imperial College London[230]) |
| ptCMV-EGFP-TRIM12 | NP_001355683.1 | 53 | *Mus musculus* | PCR from iBMDM murine cDNA, ptCMV-EGFP plasmid derived from pEGFP-N1 plasmid (Clontech) (T. Thurston Laboratory, Imperial College London[230]) |
| ptCMV-EGFP-TRIM13 | NP_005789.2 | 47 | *Homo sapiens* | PCR from THP1 human cDNA, ptCMV-EGFP plasmid derived from pEGFP-N1 plasmid (Clontech) (T. Thurston Laboratory, Imperial College London[230]) |

| | | | | |
|---|---|---|---|---|
| ptCMV-EGFP-TRIM14 | NP_055603.2 | 50 | *Homo sapiens* | PCR from mixed human cDNA, ptCMV-EGFP plasmid derived from pEGFP-N1 plasmid (Clontech) (T. Thurston Laboratory, Imperial College London[230]) |
| ptCMV-EGFP-TRIM15 | NP_150232.2 | 52 | *Homo sapiens* | PCR from mixed human cDNA, ptCMV-EGFP plasmid derived from pEGFP-N1 plasmid (Clontech) (T. Thurston Laboratory, Imperial College London[230]) |
| ptCMV-EGFP-TRIM16 | NP_001335048.1 | 64 | *Homo sapiens* | PCR from mixed human cDNA , ptCMV-EGFP plasmid derived from pEGFP-N1 plasmid (Clontech) (T. Thurston Laboratory, Imperial College London[230]) |
| ptCMV-EGFP-TRIM17 | NP_001020111.1 | 54 | *Homo sapiens* | PCR from mixed human cDNA, ptCMV-EGFP plasmid derived from pEGFP-N1 plasmid (Clontech) (T. Thurston Laboratory, Imperial College London[230]) |
| ptCMV-EGFP-TRIM18 | NP_000372.1 | 72 | *Homo sapiens* | PCR from Caco2 human cDNA, ptCMV-EGFP plasmid derived from pEGFP-N1 plasmid (Clontech) (T. Thurston Laboratory, Imperial College London[230]) |
| ptCMV-EGFP-TRIM19 | NP_150241.2 | 98 | *Homo sapiens* | PCR from mixed human cDNA, ptCMV-EGFP plasmid derived from pEGFP-N1 plasmid (Clontech) (T. Thurston Laboratory, Imperial College London[230]) |
| ptCMV-EGFP-TRIM20 | NP_000234.1 | 86 | *Homo sapiens* | PCR from mixed human cDNA, ptCMV-EGFP plasmid derived from pEGFP-N1 plasmid (Clontech) (T. Thurston Laboratory, Imperial College London[230]) |
| ptCMV-EGFP-TRIM21 | NP_003132.2 | 54 | *Homo sapiens* | PCR from U937 human cDNA, ptCMV-EGFP plasmid derived from pEGFP-N1 plasmid (Clontech) (T. Thurston Laboratory, Imperial College London[230]) |
| ptCMV-EGFP-TRIM22 | NP_006065.2 | 57 | *Homo sapiens* | Subcloned from M6P-EGFP vector generated by PCR of U937 human cDNA, ptCMV-EGFP plasmid derived from pEGFP-N1 plasmid (Clontech) (T. Thurston Laboratory, Imperial College London[230]) |
| ptCMV-EGFP-TRIM23 | NP_001647.1 | 64 | *Homo sapiens* | Subcloned from plasmid gifted from Michaela Gack[27] (T. Thurston Laboratory, Imperial College London) |
| ptCMV-EGFP-TRIM24 | NP_056989.2 | 117 | *Homo sapiens* | PCR from plasmid synthesised by Open Biosystems, ptCMV-EGFP plasmid derived from pEGFP-N1 plasmid (Clontech) (T. Thurston |

| | | | | |
|---|---|---|---|---|
| | | | | Laboratory, Imperial College London[230]) |
| ptCMV-EGFP-TRIM25 | NP_005073.2 | 71 | *Homo sapiens* | PCR from mixed human cDNA, ptCMV-EGFP plasmid derived from pEGFP-N1 plasmid (Clontech) (T. Thurston Laboratory, Imperial College London[230]) |
| ptCMV-EGFP-TRIM26 | NP_001229712.1 | 62 | *Homo sapiens* | PCR from U937 human cDNA, ptCMV-EGFP plasmid derived from pEGFP-N1 plasmid (Clontech) (T. Thurston Laboratory, Imperial College London[230]) |
| ptCMV-EGFP-TRIM27 | NP_006501.1 | 58 | *Homo sapiens* | PCR from 293ET human cDNA, ptCMV-EGFP plasmid derived from pEGFP-N1 plasmid (Clontech) (T. Thurston Laboratory, Imperial College London[230]) |
| ptCMV-EGFP-TRIM28 | NP_005753.1 | 100 | *Homo sapiens* | PCR from mixed human cDNA, ptCMV-EGFP plasmid derived from pEGFP-N1 plasmid (Clontech) (T. Thurston Laboratory, Imperial College London[230]) |
| ptCMV-EGFP-TRIM29 | NP_036233.2 | 66 | *Homo sapiens* | PCR from mixed human cDNA, ptCMV-EGFP plasmid derived from pEGFP-N1 plasmid (Clontech) (T. Thurston Laboratory, Imperial College London[230]) |
| ptCMV-EGFP-TRIM30 | NP_001344396.1 | 105 | *Mus musculus* | Subcloned from M6P-EGFP vector generated by PCR of iBMDM murine cDNA, ptCMV-EGFP plasmid derived from pEGFP-N1 plasmid (Clontech) (T. Thurston Laboratory, Imperial College London[230]) |
| ptCMV-EGFP-TRIM31 | NP_008959.3 | 48 | *Homo sapiens* | PCR from mixed human cDNA, ptCMV-EGFP plasmid derived from pEGFP-N1 plasmid (Clontech) (T. Thurston Laboratory, Imperial College London[230]) |
| ptCMV-EGFP-TRIM32 | NP_001093149.1 | 72 | *Homo sapiens* | PCR from mixed human cDNA, ptCMV-EGFP plasmid derived from pEGFP-N1 plasmid (Clontech) (T. Thurston Laboratory, Imperial College London[230]) |
| ptCMV-EGFP-TRIM33 | NP_056990.3 | 150 | *Homo sapiens* | PCR from mixed human cDNA, ptCMV-EGFP plasmid derived from pEGFP-N1 plasmid (Clontech) (T. Thurston Laboratory, Imperial College London[230]) |
| ptCMV-EGFP-TRIM34 | NP_001003827.1 | 36 | *Homo sapiens* | PCR from mixed human cDNA, ptCMV-EGFP plasmid derived from pEGFP-N1 plasmid (Clontech) (T. Thurston Laboratory, Imperial College London[230]) |
| ptCMV-EGFP-TRIM35 | NP_741983.2 | 55 | *Homo sapiens* | PCR from mixed human cDNA, ptCMV-EGFP |

| | | | | |
|---|---|---|---|---|
| | | | | plasmid derived from pEGFP-N1 plasmid (Clontech) (T. Thurston Laboratory, Imperial College London[230]) |
| ptCMV-EGFP-TRIM36 | NP_061170.2 | 83 | *Homo sapiens* | PCR from mixed human cDNA, ptCMV-EGFP plasmid derived from pEGFP-N1 plasmid (Clontech) (T. Thurston Laboratory, Imperial College London[230]) |
| ptCMV-EGFP-TRIM37 | NP_001005207.1 | 130 | *Homo sapiens* | PCR from mixed human cDNA, ptCMV-EGFP plasmid derived from pEGFP-N1 plasmid (Clontech) (T. Thurston Laboratory, Imperial College London[230]) |
| ptCMV-EGFP-TRIM38 | NP_006346.1 | 53 | *Homo sapiens* | PCR from mixed human cDNA, ptCMV-EGFP plasmid derived from pEGFP-N1 plasmid (Clontech) (T. Thurston Laboratory, Imperial College London[230]) |
| ptCMV-EGFP-TRIM39 | NP_067076.2 | 60 | *Homo sapiens* | PCR from mixed human cDNA, ptCMV-EGFP plasmid derived from pEGFP-N1 plasmid (Clontech) (T. Thurston Laboratory, Imperial College London[230]) |
| ptCMV-EGFP-TRIM40 | NP_001273562.1 | 29 | *Homo sapiens* | PCR from mixed human cDNA, ptCMV-EGFP plasmid derived from pEGFP-N1 plasmid (Clontech) (T. Thurston Laboratory, Imperial College London[230]) |
| ptCMV-EGFP-TRIM41 | NP_291027.3 | 72 | *Homo sapiens* | PCR from mixed human cDNA, ptCMV-EGFP plasmid derived from pEGFP-N1 plasmid (Clontech) (T. Thurston Laboratory, Imperial College London[230]) |
| ptCMV-EGFP-TRIM42 | NP_689829.3 | 80 | *Homo sapiens* | PCR from mixed human cDNA, ptCMV-EGFP plasmid derived from pEGFP-N1 plasmid (Clontech) (T. Thurston Laboratory, Imperial College London[230]) |
| ptCMV-EGFP-TRIM43 | NP_620155.1 | 52 | *Homo sapiens* | PCR from mixed human cDNA, ptCMV-EGFP plasmid derived from pEGFP-N1 plasmid (Clontech) (T. Thurston Laboratory, Imperial College London[230]) |
| ptCMV-EGFP-TRIM44 | NP_060053.2 | 38 | *Homo sapiens* | PCR from mixed human cDNA, ptCMV-EGFP plasmid derived from pEGFP-N1 plasmid (Clontech) (T. Thurston Laboratory, Imperial College London[230]) |
| ptCMV-EGFP-TRIM45 | NP_079464.2 | 62 | *Homo sapiens* | PCR from mixed human cDNA, ptCMV-EGFP plasmid derived from pEGFP-N1 plasmid (Clontech) (T. Thurston Laboratory, Imperial College London[230]) |

| | | | | |
|---|---|---|---|---|
| ptCMV-EGFP-TRIM46 | NP_079334.3 | 83 | *Homo sapiens* | Subcloned from M6P-EGFP vector generated by PCR of mixed human cDNA, ptCMV-EGFP plasmid derived from pEGFP-N1 plasmid (Clontech) (T. Thurston Laboratory, Imperial College London[230]) |
| ptCMV-EGFP-TRIM47 | NP_258411.2 | 70 | *Homo sapiens* | PCR from mixed human cDNA, ptCMV-EGFP plasmid derived from pEGFP-N1 plasmid (Clontech) (T. Thurston Laboratory, Imperial College London[230]) |
| ptCMV-EGFP-TRIM48 | NP_077019.2 | 24 | *Homo sapiens* | PCR from mixed human cDNA, ptCMV-EGFP plasmid derived from pEGFP-N1 plasmid (Clontech) (T. Thurston Laboratory, Imperial College London[230]) |
| ptCMV-EGFP-TRIM49 | NP_065091.1 | 53 | *Homo sapiens* | PCR from mixed human cDNA, ptCMV-EGFP plasmid derived from pEGFP-N1 plasmid (Clontech) (T. Thurston Laboratory, Imperial College London[230]) |
| ptCMV-EGFP-TRIM50 | NP_001268380.1 | 55 | *Homo sapiens* | Subcloned from plasmid gifted from Michaela Gack[27], ptCMV-EGFP plasmid derived from pEGFP-N1 plasmid (Clontech) (T. Thurston Laboratory, Imperial College London[230]) |
| ptCMV-EGFP-TRIM51 | NP_116070.2 | 53 | *Homo sapiens* | PCR from mixed human cDNA, ptCMV-EGFP plasmid derived from pEGFP-N1 plasmid (Clontech) (T. Thurston Laboratory, Imperial College London[230]) |
| ptCMV-EGFP-TRIM52 | NP_116154.1 | 33 | *Homo sapiens* | PCR from mixed human cDNA, ptCMV-EGFP plasmid derived from pEGFP-N1 plasmid (Clontech) (T. Thurston Laboratory, Imperial College London[230]) |
| ptCMV-EGFP-TRIM54 | NP_115935.3 | 40 | *Homo sapiens* | PCR from plasmid synthesised by Open Biosystems, ptCMV-EGFP plasmid derived from pEGFP-N1 plasmid (Clontech) (T. Thurston Laboratory, Imperial College London[230]) |
| ptCMV-EGFP-TRIM55 | NP_908973.1 | 60 | *Homo sapiens* | PCR from plasmid synthesised by Open Biosystems, ptCMV-EGFP plasmid derived from pEGFP-N1 plasmid (Clontech) (T. Thurston Laboratory, Imperial College London[230]) |
| ptCMV-EGFP-TRIM56 | NP_112223.1 | 81 | *Homo sapiens* | PCR from mixed human cDNA, ptCMV-EGFP plasmid derived from pEGFP-N1 plasmid (Clontech) (T. Thurston Laboratory, Imperial College London[230]) |

| | | | | |
|---|---|---|---|---|
| ptCMV-EGFP-TRIM58 | NP_056246.3 | 55 | *Homo sapiens* | PCR from mixed human cDNA, ptCMV-EGFP plasmid derived from pEGFP-N1 plasmid (Clontech) (T. Thurston Laboratory, Imperial College London[230]) |
| ptCMV-EGFP-TRIM59 | NP_775107.1 | 47 | *Homo sapiens* | PCR from mixed human cDNA, ptCMV-EGFP plasmid derived from pEGFP-N1 plasmid (Clontech) (T. Thurston Laboratory, Imperial College London[230]) |
| ptCMV-EGFP-TRIM60 | NP_001244954.1 | 55 | *Homo sapiens* | Subcloned from M6P-EGFP vector generated by subcloning from plasmid synthesised by Open Biosystems, ptCMV-EGFP plasmid derived from pEGFP-N1 plasmid (Clontech) (T. Thurston Laboratory, Imperial College London[230]) |
| ptCMV-EGFP-TRIM61 | NP_001401833.1 | 24 | *Homo sapiens* | PCR from plasmid synthesised by Open Biosystems, ptCMV-EGFP plasmid derived from pEGFP-N1 plasmid (Clontech) (T. Thurston Laboratory, Imperial College London[230]) |
| ptCMV-EGFP-TRIM62 | NP_060677.2 | 54 | *Homo sapiens* | PCR from plasmid synthesised by Open Biosystems, ptCMV-EGFP plasmid derived from pEGFP-N1 plasmid (Clontech) (T. Thurston Laboratory, Imperial College London[230]) |
| ptCMV-EGFP-TRIM63 | NP_115977.2 | 40 | *Homo sapiens* | PCR from mixed human cDNA, ptCMV-EGFP plasmid derived from pEGFP-N1 plasmid (Clontech) (T. Thurston Laboratory, Imperial College London[230]) |
| ptCMV-EGFP-TRIM64 | NP_001129958.1 | 52 | *Homo sapiens* | PCR from mixed human cDNA, ptCMV-EGFP plasmid derived from pEGFP-N1 plasmid (Clontech) (T. Thurston Laboratory, Imperial College London[230]) |
| ptCMV-EGFP-TRIM65 | NP_775818.2 | 57 | *Homo sapiens* | Subcloned from GenScript synthesised pcDNA-GFP vector, ptCMV-EGFP plasmid derived from pEGFP-N1 plasmid (Clontech) (T. Thurston Laboratory, Imperial College London[230]) |
| ptCMV-EGFP-TRIM66 | NP_001374951.1 | 135 | *Homo sapiens* | PCR from mixed human cDNA, ptCMV-EGFP plasmid derived from pEGFP-N1 plasmid (Clontech) (T. Thurston Laboratory, Imperial College London[230]) |
| ptCMV-EGFP-TRIM67 | NP_001004342.3 | 84 | *Homo sapiens* | Subcloned from GenScript synthesised pcDNA-GFP vector, ptCMV-EGFP plasmid derived from pEGFP-N1 plasmid (Clontech) (T. Thurston |

| | | | | |
|---|---|---|---|---|
| | | | | Laboratory, Imperial College London[230]) |
| ptCMV-EGFP-TRIM68 | NP_060543.5 | 56 | *Homo sapiens* | PCR from mixed human cDNA, ptCMV-EGFP plasmid derived from pEGFP-N1 plasmid (Clontech) (T. Thurston Laboratory, Imperial College London[230]) |
| ptCMV-EGFP-TRIM69 | NP_892030.3 | 57 | *Homo sapiens* | PCR from mixed human cDNA, ptCMV-EGFP plasmid derived from pEGFP-N1 plasmid (Clontech) (T. Thurston Laboratory, Imperial College London[230]) |
| ptCMV-EGFP-TRIM70 | NP_001340153.1 | 40 | *Homo sapiens* | PCR from mixed human cDNA, ptCMV-EGFP plasmid derived from pEGFP-N1 plasmid (Clontech) (T. Thurston Laboratory, Imperial College London[230]) |
| ptCMV-EGFP-TRIM71 | NP_001034200.1 | 93 | *Homo sapiens* | PCR from mixed human cDNA, ptCMV-EGFP plasmid derived from pEGFP-N1 plasmid (Clontech) (T. Thurston Laboratory, Imperial College London[230]) |
| ptCMV-EGFP-TRIM72 | NP_001008275.2 | 53 | *Homo sapiens* | Subcloned from M6P-EGFP plasmid synthesised by Adolfo García-Sastre, , ptCMV-EGFP plasmid derived from pEGFP-N1 plasmid (Clontech) (T. Thurston Laboratory, Imperial College London[230]) |
| ptCMV-EGFP-TRIM73 | NP_944606.2 | 29 | *Homo sapiens* | PCR from mixed human cDNA, ptCMV-EGFP plasmid derived from pEGFP-N1 plasmid (Clontech) (T. Thurston Laboratory, Imperial College London[230]) |
| ptCMV-EGFP-TRIM74 | NP_001304744.1 | 29 | *Homo sapiens* | PCR from mixed human cDNA, ptCMV-EGFP plasmid derived from pEGFP-N1 plasmid (Clontech) (T. Thurston Laboratory, Imperial College London[230]) |
| ptCMV-EGFP-TRIM75 | NP_001382999.1 | 54 | *Homo sapiens* | Subcloned from M6P-EGFP vector generated by PCR of mixed human cDNA, ptCMV-EGFP plasmid derived from pEGFP-N1 plasmid (Clontech) (T. Thurston Laboratory, Imperial College London[230]) |
| ptCMV-EGFP-TRIM77 | NP_001139634.1 | 52 | *Homo sapiens* | Subcloned from M6P-EGFP vector generated by PCR of mixed human cDNA, ptCMV-EGFP plasmid derived from pEGFP-N1 plasmid (Clontech) (T. Thurston Laboratory, Imperial College London[230]) |
| ptCMV-EGFP-TRIML1 | NP_848651.2 | 53 | *Homo sapiens* | Subcloned from M6P-EGFP vector generated by PCR of mixed human cDNA, ptCMV-EGFP plasmid derived from pEGFP-N1 plasmid (Clontech) (T. |

| | | | | |
|---|---|---|---|---|
| | | | | Thurston Laboratory, Imperial College London[230]) |
| ptCMV-EGFP-TRIML2 | NP_001290348.1 | 44 | *Homo sapiens* | PCR from mixed human cDNA, ptCMV-EGFP plasmid derived from pEGFP-N1 plasmid (Clontech) (T. Thurston Laboratory, Imperial College London[230]) |
